# Modelling The Brain’s Response To Natural Scenes In The Bottleneck Space

**DOI:** 10.1101/2023.07.30.551149

**Authors:** Henry Ndubuaku

## Abstract

Computational models that mirror the brain’s behaviour can help us understand human intelligence and SOTA techniques for modelling the brain’s response to visual stimuli use deep neural networks. However, the best-performing vision models are compute-intensive and functional brain activities are represented by high-dimensional matrices which exacerbate this inefficiency. To this end, we propose a novel approach which showed significant performance improvements by 1) Projecting both the visual stimuli features and brain responses to low-dimensional vectors and using a non-linear neural network to learn the mapping in the latent space. 2) Simultaneously modelling all vertices in the visual cortices of both the left and right hemispheres using an objective we call “Racing Loss”. 3) Incorporating tiny leaks of the ground truth during training of this network. 4) First pre-training this network on all subjects then fine-tuning on each. We show that our method additionally achieved 12% higher Noise-Normalized Mean Correlation Scores compared to fully fine-tuning large vision models to the high-dimensional brain responses.

## 1 Introduction

The superiority of deep neural networks in modelling the brain’s response to visual stimuli has been demonstrated in various works such as [1], and [2]. These studies have shown that neural networks can effectively capture complex visual representations, and their performance improves with scale, as indicated by [3]. For problems involving predicting the brain’s response to natural scenes, [5] demonstrated that voxel-wise encoding models, enhanced by integrating features from pre-trained neural networks and cortical network information, achieved the best performance. These techniques show promise in learning the brain’s response to visual stimuli of natural scenes and understanding our brains.

Despite these advancements, Image-to-fMRI translation is computationally expensive. The resource-intensive nature of large-scale models limits their practicality in certain contexts. Also, the brain’s functional response to stimuli is typically represented by Functional Magnetic Resonance Imaging (fMRI). The fMRI for the visual cortices is typically represented by matrices with 10s of thousands of elements per hemisphere. This study, therefore, focused on exploring more efficient approaches for learning the brain’s response to visual stimuli from natural scenes. The Natural Scenes Dataset (NSD)[6] was utilized for this study. As formally described in the paper [6], this consists of brain responses from 8 human participants to in total 73,000 different visual scenes. The brain responses were measured with functional MRI (fMRI). The scenes in the dataset were selected to be diverse and representative of natural scenes that humans typically encounter in their daily lives.

## 2 Method

We approached the problem with multiple beliefs. Firstly, we considered the benefits of learning a lower-dimensional linear transform of the fMRI data. By doing so, we aimed to enable the neural network to efficiently learn the fMRI mapping itself while allowing for flexibility in configuring this low-dimension neural network. Secondly, we aimed to promote more effective cross-hemispheric learning and a neural network with low-dimension inputs and targets gives room to simultaneously learn the mappings of both the left and right visual cortex without performance trade-offs. This work introduces a loss function which ensures a balance between the left and right hemispheres during training. Thirdly, we introduced a novel training technique involving leaking tiny droplets of the fMRI ground truths during the training process. This approach, which is a form of professor forcing [4], should provide additional guidance to the neural network during training. By intermittently incorporating actual fMRI data as a reference, we aimed to enhance the model’s ability to learn meaningful representations from the visual stimuli and the corresponding brain responses. Lastly, by pre-training the model on every subject in the dataset, the network ought to learn better intermediate representations of images. This pre-training step should serve as a form of transfer learning, as the model learns global patterns from a broad range of individuals, which subsequently boosts its performance when trained on individual subjects. In the following sections, we provided detailed descriptions of our methods and presented the results of our experiments. We demonstrated the effectiveness of each approach in improving the ability of a low-dimension neural network to map accurately map image stimuli to visual cortex activity.

### 2.1 Feature Extraction

Our image features were obtained from the last hidden layer of a pre-trained CLIP Vision Transformer Base 32 [9] for the following reasons. First, transfer learning is crucial for image-to-fMRI translation due to the scarcity of labelled fMRI data compared to abundant image datasets. Second, humans reason primarily reason in language [12], this vision model is trained to produce embeddings of images that cluster with the embeddings of its equivalent linguistic description in a vector space. To mine more multimodal relations, we tried extracting describing captions for each image stimuli using BLIP [10], and concatenating its embeddings to that of the images. This however failed to yield tangible improvements enough to justify its resource consumption. The obtained captions were not descriptive enough to emulate reasoning over an image, descriptiveness has been proven to improve performance in multimodal learning. [11].

### 2.2 Learning In The Bottleneck Space

The bottleneck can be seen as a vector space where data is compressed to a lower dimension while retaining its information. Principal Component Analysis (PCA) allows for reducing dimensionality while preserving as much information as possible [13]. In our setup, PCA is used to not only project the image features to a lower dimension vector (300, down from 37,800) as prevalent in works like [14], we additionally transform the ground truth fMRI into lower dimension vectors (75 each for the left and right hemispheres, down from about 20k). The whole training then occurs in the bottleneck space as shown in 2.2. At inference, the predicted low-dimension FMRI is inversely transformed back to the high-dimension space. With this, the bottleneck network 2.2 can be scaled without exploding the number of parameters or using more resources.

**Figure 1:**
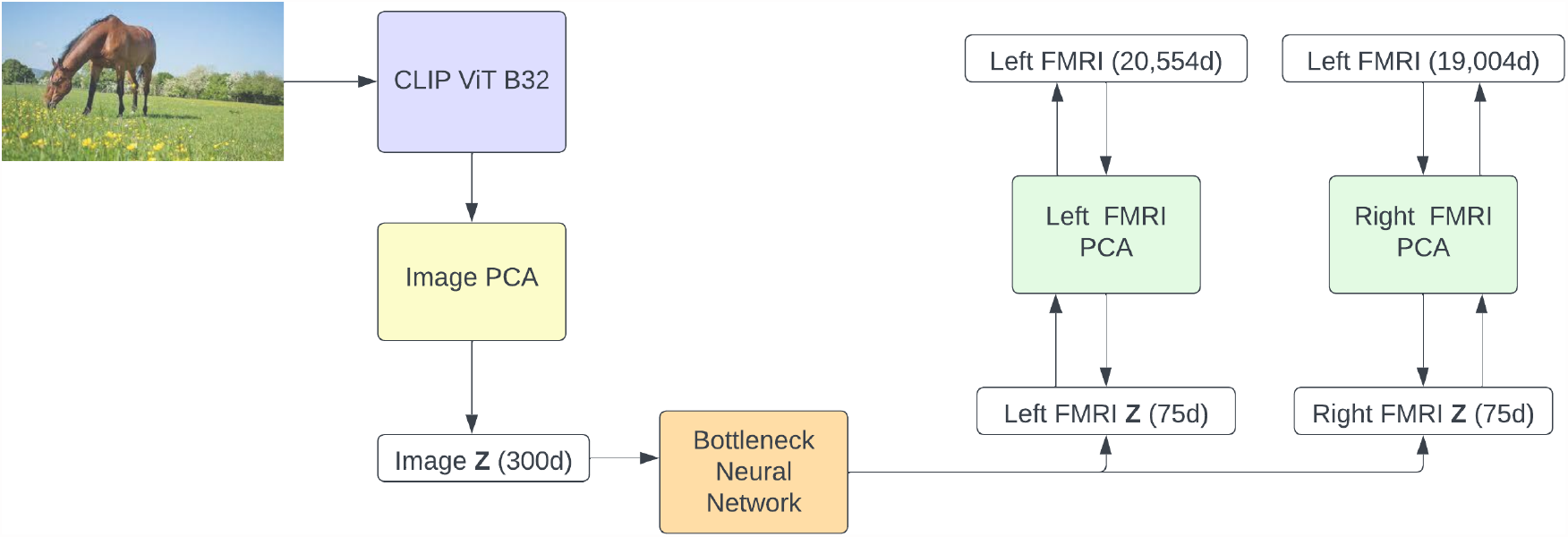
Bottleneck Learning.

**Figure 2:**
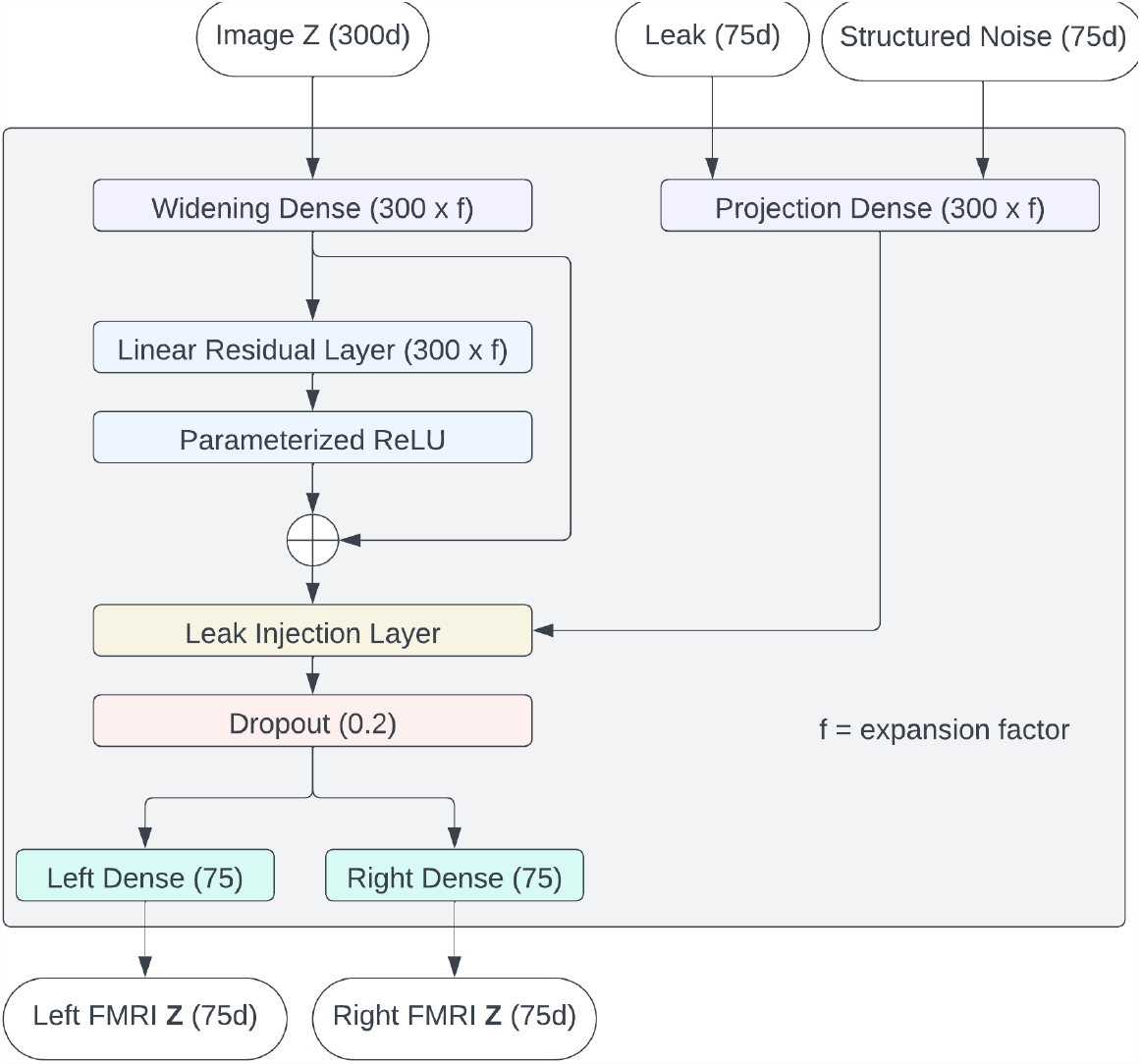
Bottleneck Neural Network.

### 2.3 Information Leaking

Considering that the goal is to learn representations of visual stimuli that align with equivalent representations in the human brain, this is a form of multimodal alignment representation-learning problem. Contrastive learning methods have proven to yield excellent results in such tasks [8]. But this is a unique case, only the artificial neural network would be learning, and the brain under study would not. This limited the benefits of contrastive learning. Drawing inspiration from professor forcing [4] and to still enjoy the advantages of learning from the fMRI, during training, the ground truth fMRI (**y**) is injected as shown in 1.

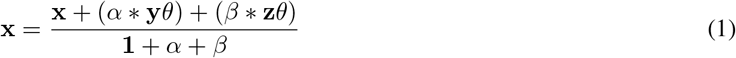

**z** is a structured noise obtained by scaling normally sampled eigenvectors by the square root of the eigenvalues of the top n principal components selected when down-sampling the fMRI using PCA. **y** and **z** are parameterised by the same projection layer *theta. alpha* and *beta* control the ratio of **y** to **z** 3. The denominator scales the result of adding terms in the numerator down.

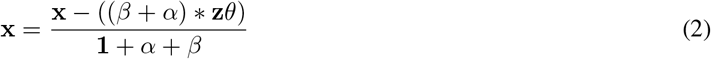

During inference, the ground truth is unavailable and as such the noise is injected as shown in 2. Removing the noise and scaling yielded better results compared to adding only the structured noise **z**.

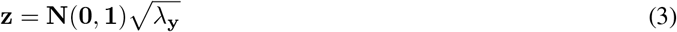

### 2.4 Racing Loss

On training a single neural network to map both the left and right visual cortices, we observed that the network often learned to gravitate towards either side. To sensor balance, we propose using “Racing Loss” 4. To gain an intuitive understanding of this loss function, consider a human with two legs moving from a starting position to an endpoint. One could say, both legs are racing towards the endpoint and the human maintains balance by pushing the lagging leg forward. In a similar fashion, for each update, the loss from the lagging cortex is weighted more.

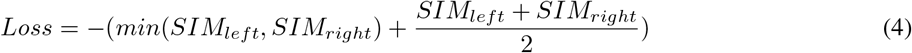

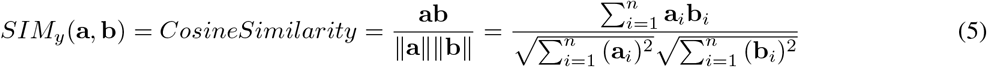

### 2.5 Group Pre-training

Merging brain information from multiple candidates can be accomplished through a technique called hyperalignment[15]. The basic idea behind hyperalignment is to treat cortical patterns as vectors corresponding to locations in a high dimensional space, where each axis reflects a measurement of that pattern (e.g., voxel activity). The algorithm then finds a transformation that maps the vectors from each subject into the same common information space. This transformation is typically found using a technique called Procrustes analysis, which minimizes the difference between the transformed vectors from different subjects.

Building on this idea, we initially pre-trained a backbone bottleneck neural network on data from all the candidates, using their aligned brain responses. This pre-training step was aimed at creating more robust initial weights for the network, enabling it to capture more general and informative features from the combined data, hyperalignemt facilitated combining FMRI of various subjects. Subsequently, we fine-tuned the network for each individual subject. By doing this, the network can adapt to each subject’s unique brain characteristics while still benefiting from the knowledge gained during the initial pre-training step.

## 3 Experimentation and Results

Firstly, to assess the encoding accuracy of our technique in predicting brain responses, evaluation was carried out on the testing set withheld as part of the Algonaut Challenge 2023 [7]. This involved determining the mean noise-normalized encoding accuracy across all vertices, considering all subjects and hemispheres, details can be found in the paper.

The baseline model was a CLIP-ViT-B32 fine-tuned separately for each of the 8 subjects while saving the best-performing weights. This baseline was trained to map visual stimuli to their corresponding brain activities (via FMRI) in their original dimensions. The Mean Squared Error (MSE) was utilised.

In our setup (referenced as “F-BNet”), the weights of the CLIP-ViT-B32 are frozen to ensure the learning process only occurs in the bottleneck space. The average training time for the baseline model on a single Nvidia A40 GPU was 6 minutes and 43 seconds in 10 epochs, while the F-BNet model took only 43 seconds for the same training process, both using a batch size of 32. We tested 3 more variants, F-Net without leak injection, F-Net with MSE in place of racing loss, and F-Net with random weight initialization. Results are summarized in 1, where each column indicates the subject number and hemisphere (left or right).

**Table 1:** Mean noise-normalized encoding accuracy across all the vertices of all subjects and hemispheres.

| Vertices | All | 1L | 1R | 2L | 2R | 3L | 3R | 4L | 4R | 5L | 5R | 6L | 6R | 7L | 7R | 8L | 8R |
| --- | --- | --- | --- | --- | --- | --- | --- | --- | --- | --- | --- | --- | --- | --- | --- | --- | --- |
| CLIP-ViT-B32 | 41.41 | 39.57 | 39.10 | 35.40 | 35.70 | 48.97 | 47.89 | 39.71 | 42.62 | 42.04 | 42.49 | 43.87 | 46.29 | 31.78 | 33.68 | 46.25 | 47.37 |
| F-BNet | 46.27 | 47.01 | 46.21 | 42.06 | 42.07 | 50.57 | 50.01 | 44.91 | 48.46 | 44.44 | 45.02 | 50.54 | 51.14 | 37.43 | 37.66 | 51.17 | 51.20 |
| F-BNet (No Pretrain) | 44.86 | 45.87 | 45.21 | 40.43 | 40.97 | 48.76 | 48.44 | 44.14 | 47.66 | 43.94 | 44.33 | 48.46 | 49.86 | 36.18 | 34.90 | 48.14 | 50.57 |
| F-BNet (No Leak) | 45.54 | 45.87 | 44.71 | 41.01 | 41.70 | 49.28 | 49.89 | 44.83 | 47.75 | 44.74 | 44.57 | 50.07 | 50.09 | 37.50 | 36.82 | 48.32 | 51.54 |
| F-BNet (MSE) | 44.34 | 44.04 | 44.62 | 43.18 | 43.78 | 36.08 | 37.22 | 36.04 | 38.98 | 43.33 | 42.14 | 33.72 | 36.97 | 35.37 | 35.51 | 29.79 | 30.56 |

## 4 Discussion

The remarkable reduction in memory usage by the F-BNet model allowed for a significant increase in batch size, which could further shrink training time to approximately 15.6 seconds. This highlights the efficiency and advantage of F-BNet in terms of memory consumption, enabling faster training by handling larger batch sizes more effectively compared to the baseline model.

It is worth noting that PCA is a lossy compression technique, hence this inverse transform does not reconstruct perfectly. We found the reconstruction performance to increase with more principal components while the neural networks in reverse perform better as components reduce. Error increasing in either direction weakened F-BNet’s performance, we recommend finding the perfect learning-reconstruction error balance for each dataset. 75 principal components were ideal for the Natural Scenes Dataset. Again, the expansion factor for the widening layer saturates at 20 for 75 principal components. Finally, the alpha/beta ratio in the leak injection layer performs best at 10:1 and we feel the leak injection approach during inference needs more investigation. While racing loss improved performance, more investigation is required to understand its effectiveness.

Our approach is neither expected to be the best nor the most neuromorphic approach for computationally generating fMRI representations of visual stimuli. We instead propose fundamental concepts to facilitate working with lower dimensional projections of both the input and output variables, hence better computation efficiency. To our surprise, these concepts in synergy can improve encoding accuracy if properly tuned. We provide the source codes for reproducing the experiment as well as the full results on various brain regions of interest in our repository here.

## 5 Conclusion and Future Work

In conclusion, our study explored ways to efficiently model how the brain responds to visual stimuli. We introduced a novel approach that projects images and brain responses to low-dimensional vectors and learns non-linear mapping. To enhance learning, we cleverly injected small portions of ground truth brain data during training, while during inference, we used controlled noise injection. We also used a novel loss function to ensure balanced learning between the brain’s left and right visual hemispheres. Finally, we employed a group pertaining on the hyper-aligned brain responses of all subjects, then fine-tuned on each subject.

The results were promising, our method proved to be more practical and resource-efficient. It outperformed the baseline model, achieving 12% higher Noise-Normalized Mean Correlation Scores. We believe each concept in our approach has the potential for broader applications beyond this specific dataset and domain. Future works could explore each in depth.

## Supporting information

Source code & results.

## Notes

https://github.com/HMUNACHI/FBNET

### Competing Interest Statement

The authors have declared no competing interest.

### Summary of Updates

A discussion section was added and grammatical errors were corrected.

