## Supplementary material for "Modelling The Brain’s Response To Natural Scenes In The Bottleneck Space": Source code & results.: extraction_scripts.py

```
import os
import pathlib
from PIL import Image
import jax.numpy as jnp
from transformers import AutoProcessor, FlaxCLIPVisionModel

""""
Note: This is customised for how the dataset was structured for ease of use
"""

clip = FlaxCLIPVisionModel.from_pretrained("openai/clip-vit-base-patch32")
clip_processor = AutoProcessor.from_pretrained("openai/clip-vit-base-patch32")

for subj in range(1,9):

    subj = str(subj)
    print("Extracting Features from Subject: {}".format(subj))
    path = "data/subj0" + subj

    train_img_dir  = os.path.join('NSD_dataset', 'subj0'+subj, 'training_split', 'training_images')
    train_img_list = list(pathlib.Path(train_img_dir).glob('*png'))
    train_img_list.sort()
    train_images = jnp.array(list(map(Image.open, train_img_list)))
    jnp.save(path + "/train_images.npy", train_images)
    del train_img_list

    batch_size = 10
    start, end = 0, batch_size
    size = len(train_images) // batch_size * 10
    txts, imgs = [], []

    while end <= size:
        img = train_images[start:end]
        inputs = clip_processor(images=img,return_tensors="np")
        outputs = clip(**inputs)
        imgs.extend(outputs.last_hidden_state)
        print(end)
        start = end
        end += batch_size

    imgs = jnp.array(imgs)
    imgs = imgs.reshape((imgs.shape[0],-1))
    jnp.save(path + "/train_X.npy", imgs)


    test_img_dir  = os.path.join('NSD_dataset', 'subj0'+subj, 'test_split', 'test_images')
    test_img_list = list(pathlib.Path(test_img_dir).glob('*png'))
    test_img_list.sort()
    test_images = jnp.array(list(map(Image.open, test_img_list)))

    inputs = clip_processor(images=test_images,return_tensors="np")
    outputs = clip(**inputs)
    imgs = outputs.last_hidden_state
    imgs = imgs.reshape((imgs.shape[0],-1))
    jnp.save(path + "/test_X.npy", imgs)
```
