## Supplementary material for "Modelling The Brain’s Response To Natural Scenes In The Bottleneck Space": Source code & results.: extraction_utils.py

```
import jax
import numpy as np
import jax.numpy as jnp

def zero_pad_sequence_3d(array, last_dim):
    """
    """
    new_array = np.zeros((len(array),21,last_dim),dtype=array[0].dtype)
    for idx, x in enumerate(array):
        mid_dim = min(x.shape[0], 21)
        new_array[idx, :mid_dim, :] = x[:mid_dim, :]
    return new_array

def zero_pad_sequence_2d(array, last_dim):
    """
    """
    g = []
    for x in array:
        y = np.zeros((len(x),last_dim),dtype=x.dtype)
        print(y.shape, y[:, :len(x[0])].shape, x.shape)
        y[:, :len(x[0])] = x
        g.append(y)
    return np.concatenate(g, axis=0)

@jax.vmap
def get_flattened_svd(image):
    """
    """
    U, S, Vh = jnp.linalg.svd(image, full_matrices=False)
    S = jnp.diag(S)
    y = jnp.dot(S * U, Vh)
    return y.reshape(-1)
```
