## Supplementary material for "Modelling The Brain’s Response To Natural Scenes In The Bottleneck Space": Source code & results.: environment.yml

```
name: jax
channels:
  - conda-forge
  - defaults
dependencies:
  - _libgcc_mutex=0.1=main
  - _openmp_mutex=5.1=1_gnu
  - asttokens=2.2.1=pyhd8ed1ab_0
  - backcall=0.2.0=pyh9f0ad1d_0
  - backports=1.0=pyhd8ed1ab_3
  - backports.functools_lru_cache=1.6.4=pyhd8ed1ab_0
  - bzip2=1.0.8=h7b6447c_0
  - ca-certificates=2023.5.7=hbcca054_0
  - cudatoolkit=11.8.0=h6a678d5_0
  - debugpy=1.5.1=py310h295c915_0
  - decorator=5.1.1=pyhd8ed1ab_0
  - entrypoints=0.4=pyhd8ed1ab_0
  - executing=1.2.0=pyhd8ed1ab_0
  - ipykernel=6.15.0=pyh210e3f2_0
  - ipython=8.14.0=pyh41d4057_0
  - jedi=0.18.2=pyhd8ed1ab_0
  - jupyter_client=7.3.4=pyhd8ed1ab_0
  - jupyter_core=5.3.1=py310hff52083_0
  - ld_impl_linux-64=2.38=h1181459_1
  - libffi=3.3=he6710b0_2
  - libgcc-ng=11.2.0=h1234567_1
  - libgomp=11.2.0=h1234567_1
  - libsodium=1.0.18=h36c2ea0_1
  - libstdcxx-ng=11.2.0=h1234567_1
  - libuuid=1.41.5=h5eee18b_0
  - matplotlib-inline=0.1.6=pyhd8ed1ab_0
  - ncurses=6.4=h6a678d5_0
  - nest-asyncio=1.5.6=pyhd8ed1ab_0
  - openssl=1.1.1u=h7f8727e_0
  - packaging=23.1=pyhd8ed1ab_0
  - parso=0.8.3=pyhd8ed1ab_0
  - pexpect=4.8.0=pyh1a96a4e_2
  - pickleshare=0.7.5=py_1003
  - pip=23.1.2=py310h06a4308_0
  - platformdirs=3.6.0=pyhd8ed1ab_0
  - prompt-toolkit=3.0.38=pyha770c72_0
  - prompt_toolkit=3.0.38=hd8ed1ab_0
  - psutil=5.9.0=py310h5eee18b_0
  - ptyprocess=0.7.0=pyhd3deb0d_0
  - pure_eval=0.2.2=pyhd8ed1ab_0
  - pygments=2.15.1=pyhd8ed1ab_0
  - python=3.10.0=h12debd9_5
  - python-dateutil=2.8.2=pyhd8ed1ab_0
  - python_abi=3.10=2_cp310
  - pyzmq=25.1.0=py310h6a678d5_0
  - readline=8.2=h5eee18b_0
  - setuptools=67.8.0=py310h06a4308_0
  - six=1.16.0=pyh6c4a22f_0
  - sqlite=3.41.2=h5eee18b_0
  - stack_data=0.6.2=pyhd8ed1ab_0
  - tk=8.6.12=h1ccaba5_0
  - tornado=6.1=py310h5764c6d_3
  - traitlets=5.9.0=pyhd8ed1ab_0
  - typing-extensions=4.6.3=hd8ed1ab_0
  - typing_extensions=4.6.3=pyha770c72_0
  - wcwidth=0.2.6=pyhd8ed1ab_0
  - wheel=0.38.4=py310h06a4308_0
  - xz=5.4.2=h5eee18b_0
  - zeromq=4.3.4=h9c3ff4c_1
  - zlib=1.2.13=h5eee18b_0
  - pip:
      - absl-py==1.4.0
      - cached-property==1.5.2
      - certifi==2023.5.7
      - charset-normalizer==3.1.0
      - chex==0.1.7
      - contourpy==1.1.0
      - cycler==0.11.0
      - dataclasses==0.6
      - dm-haiku==0.0.9
      - dm-tree==0.1.8
      - etils==1.3.0
      - filelock==3.12.2
      - flax==0.6.11
      - flaxmodels==0.1.2
      - fonttools==4.40.0
      - fsspec==2023.6.0
      - h5py==3.9.0
      - huggingface-hub==0.15.1
      - idna==3.4
      - importlib-resources==5.12.0
      - jax==0.4.12
      - jax-dataloader==0.0.5
      - jaxlib==0.4.12+cuda11.cudnn86
      - jmp==0.0.4
      - joblib==1.2.0
      - kiwisolver==1.4.4
      - markdown-it-py==3.0.0
      - matplotlib==3.7.1
      - mdurl==0.1.2
      - ml-dtypes==0.2.0
      - msgpack==1.0.5
      - numpy==1.25.0
      - nvidia-cublas-cu11==2022.4.8
      - nvidia-cublas-cu117==11.10.1.25
      - nvidia-cudnn-cu11==8.6.0.163
      - opt-einsum==3.3.0
      - optax==0.1.5
      - orbax-checkpoint==0.2.6
      - pandas==2.0.3
      - pillow==9.5.0
      - pyparsing==3.1.0
      - pytz==2023.3
      - pyyaml==6.0
      - regex==2023.6.3
      - requests==2.31.0
      - rich==13.4.2
      - safetensors==0.3.1
      - scikit-learn==1.2.2
      - scipy==1.10.1
      - tabulate==0.9.0
      - tensorstore==0.1.38
      - threadpoolctl==3.1.0
      - tokenizers==0.13.3
      - toolz==0.12.0
      - tqdm==4.65.0
      - transformers==4.30.2
      - tzdata==2023.3
      - urllib3==2.0.3
```
