## Supplementary material for "Modelling The Brain’s Response To Natural Scenes In The Bottleneck Space": Source code & results.: experimentation.ipynb

```
# %% [markdown]
First follow the steps outlined in the provided readme.

# %% [code]
import os
import jax
import optax
import pickle
import jax.numpy as jnp

from src.model import *
from src.trainer import train
from src.dataloader import DataObj
from src.metrics import batch_pearsonr
from src.evaluation import mean_roi_correlation, plot_results


@jax.value_and_grad
@jax.jit 
def train_forward(params, x, structured_noise, left_y, right_y):
    leak = jnp.concatenate((left_y, right_y), axis=-1)
    left_preds, right_preds = model.apply(params, x, structured_noise, leak)
    left_loss = optax.cosine_similarity(left_preds, left_y).mean()
    right_loss= optax.cosine_similarity(right_preds, right_y).mean()
    return -(jnp.minimum(left_loss, right_loss) + ((left_loss + right_loss)/2))

@jax.jit 
def eval_forward(params, x, structured_noise, left_y, right_y):
    left_preds, right_preds = model.apply(params, x, structured_noise)
    left_corr = batch_pearsonr(left_preds, left_y)
    right_corr = batch_pearsonr(right_preds, right_y)
    return left_corr, right_corr, left_preds, right_preds

# %% [code]
# Data Hyperparameters
subj = 1
parent_dir = "data/subj0"
ckpt_dir = 'checkpoints/' 
image_dim = 75 
low_dim = 75
batch_size = 128
validation_split = 0.05

# Model Hyperparameters
hidden_expansion = 20
dropout = 0.2
alpha = 0.02
beta = 0.002
area = None
roi = None

# Training Hyperparameters
epochs = 10
use_pretrained_weights = True
prepare_submission = True
show_plot = False

# %% [code]
# Run Experiment On Each Subject

for subj in range(1,9):
    print(subj)
    # Main: Load Data
    data = DataObj(subj=subj,
                parent_dir="data/subj0",
                batch_size=128,
                validation_split=0.05,
                low_dim=low_dim,
                area=None,
                roi=None)

    print(data.left_dim, data.right_dim)

    # Main: Intantiate model
    model = Model(
        image_dim=300,    
        hidden_expansion=20,            
        left_dim=low_dim,
        right_dim=low_dim,
        dropout=0.2,
        alpha=0.02,
        beta=0.002
        )

    # Main: Initialise model
    params = model.init(jax.random.PRNGKey(1), 
                        jnp.ones(data.input_shape), 
                        jnp.ones(data.noise_shape), 
                        jnp.ones(data.leak_shape))

    # Load pre-trained weights
    if use_pretrained_weights:
        params = pickle.load(open(os.path.join('checkpoints/', 'all'), "rb"))

    params, left_corrs, right_corrs = train(train_forward=train_forward,
                                eval_forward=eval_forward,
                                params=params,
                                optimizer=optax.lion(learning_rate=0.0001),
                                data=data,
                                epochs=10,
                                ckpt_dir='checkpoints/', 
                                prefix=str(subj))
    
    # Plot results:
    if show_plot:
        (left_roi_corr, 
        right_roi_corr, 
        roi_names) = mean_roi_correlation(parent_dir+str(subj),
                                        left_corrs,
                                        right_corrs)

        plot_results(left_roi_corr, right_roi_corr, roi_names)  

# %% [code]
# Count parameters
params = pickle.load(open(os.path.join('checkpoints/', 'all'), "rb"))
print(sum(x.size for x in jax.tree_leaves(params)))
```
