## Supplementary material for "Modelling The Brain’s Response To Natural Scenes In The Bottleneck Space": Source code & results.: README.md

# Modelling The Brain's Response To Natural Scenes In The Bottleneck Space.

# Abstract 
Computational models that mirror the brain's behaviour can help us understand human intelligence and SOTA techniques for modelling the brain's response to visual stimuli use deep neural networks. However, the best-performing vision models are compute-intensive and functional brain activities are represented by high-dimensional matrices which exacerbate this inefficiency. To this end, we propose a novel approach which showed significant performance improvements by 1) Projecting both the visual stimuli features and brain responses to low-dimensional vectors and using a non-linear neural network to learn the mapping in the latent space. 2) Simultaneously modelling all vertices in the visual cortices of both the left and right hemispheres using an objective we call "Racing Loss". 3) Incorporating tiny leaks of the ground truth during training of this network. 4) First pre-training this network on all subjects then fine-tuning on each. We show that our method additionally achieved 12% higher Noise-Normalized Mean Correlation Scores compared to fully fine-tuning large vision models to the high-dimensional brain responses.

# Authors
Henry Ndubuaku\
\
[Linkedin](https://www.linkedin.com/in/henry-ndubuaku-7b6350b8/)\
[Paper](https://www.biorxiv.org/content/10.1101/2023.07.30.551149v1)

# Details
![Alt text](/images/bottleneck.png "Diagram")
F-BNet's full pipeline.

![Alt text](/images/network.png "Diagram")
The internals of the bottleneck network.

![Alt text](/images/equations.png "Diagram")
The information leaking and racing loss function.

# Results
![Alt text](/images/scores.png "Diagram")

# Usage
* Step 1: Ensure you have Jax, Flax, Optax, Haiku and Jax-Dataoader setup, full dependenceies can be found in the provided environment.yaml file. All can be installed using "pip install <package-name>" but for GPU access, please checkout the installation guide on Jax's repository here: https://github.com/google/jax#installation

* Step 2: You can run from scratch, or download the globally pretrained weigthts from [here](https://drive.google.com/drive/folders/1iMnx18nGSaqKTVeeTGidWru2bFt6ZBK0?usp=sharing). preprocessed dataset for each subject can also be be found in the same link.

* Step 3: Place the downloaded files (checkpoints and data) in the outer FBNET folder.

* Step 4: Open the experimentation notebook to run an example usage of the base network.

# Note
* The metric reported in the paper require the witheld noise ceilings, this experimentation only use mean correlation without the noise normalisation and at such will yield lower results.

* The clip feature extraction is done separately using the provided scripts.
