## Supplementary material for "Modelling The Brain’s Response To Natural Scenes In The Bottleneck Space": Source code & results.: dataloader.py

```
import jax.numpy as jnp
import jax_dataloader as jdl
from src.downsampler import PCA

class DataObj:
    """
    """
    def __init__(self, 
                 subj, 
                 parent_dir, 
                 validation_split, 
                 batch_size, 
                 low_dim,
                 area=None, 
                 roi=None):
        
        path = parent_dir + str(subj)
        train_X = jnp.load(path + "/train_x_pca.npy")
        left_fmri = jnp.load(path + "/left.npy")[:len(train_X)]
        right_fmri = jnp.load(path + "/right.npy")[:len(train_X)]

        if area and roi:
            left_fmri, right_fmri = self.get_roi_fmri(path, 
                                                      left_fmri, 
                                                      right_fmri, 
                                                      area, 
                                                      roi)

        self.left_fmri_downsampler = PCA(dim=low_dim).fit(left_fmri)
        self.right_fmri_downsampler = PCA(dim=low_dim).fit(right_fmri)

        #left_Y = self.left_fmri_downsampler.transform(left_fmri)
        #right_Y = self.right_fmri_downsampler.transform(right_fmri)

        #kmeans_left = KMeans(n_clusters=10, random_state=0, n_init="auto").fit(left_Y.T)
        #kmeans_right = KMeans(n_clusters=10, random_state=0, n_init="auto").fit(right_Y.T)

        split_idx = int(len(train_X) * validation_split)

        self.train = jdl.DataLoader(
                jdl.ArrayDataset(train_X[split_idx:],
                        left_fmri[split_idx:],
                        right_fmri[split_idx:]),
                backend='jax', 
                batch_size=batch_size, 
                shuffle=True)

        self.val = (train_X[:split_idx],
                    left_fmri[:split_idx],
                    right_fmri[:split_idx])
        
        self.test = jnp.load(path + "/test_x_pca.npy")
        self.input_shape = train_X[:1].shape
        self.noise_shape = (batch_size, low_dim*2)
        self.leak_shape = (batch_size, low_dim*2)
        self.left_dim = left_fmri.shape[-1]
        self.right_dim = right_fmri.shape[-1]
        return 


    def get_roi_fmri(self, path, left_fmri, right_fmri, group, area=None):
        
        roi_mapping_files = {'visual':'mapping_prf-visualrois.npy', 
                            'bodies':'mapping_floc-bodies.npy',
                            'faces':'mapping_floc-faces.npy', 
                            'places':'mapping_floc-places.npy',
                            'words':'mapping_floc-words.npy', 
                            'stream':'mapping_streams.npy'}
        
        f = roi_mapping_files[group]
        roi_name_map = jnp.load(path+'/roi_masks/'+f, allow_pickle=True).item()
        roi_name_map = {v:k for k,v in roi_name_map.items()}
        print(roi_name_map.keys())
        

        left_roi_files = {'visual':'lh.prf-visualrois_challenge_space.npy', 
                        'bodies':'lh.floc-bodies_challenge_space.npy',
                        'faces':'lh.floc-faces_challenge_space.npy', 
                        'places':'lh.floc-places_challenge_space.npy',
                        'words':'lh.floc-words_challenge_space.npy', 
                        'stream':'lh.streams_challenge_space.npy'}

        right_roi_files = {'visual':'rh.prf-visualrois_challenge_space.npy', 
                        'bodies':'rh.floc-bodies_challenge_space.npy',
                        'faces':'rh.floc-faces_challenge_space.npy', 
                        'places':'rh.floc-places_challenge_space.npy',
                        'words':'rh.floc-words_challenge_space.npy', 
                        'stream':'rh.streams_challenge_space.npy'}
        
        left_f = left_roi_files[group]
        left_rois = jnp.load(path+'/roi_masks/'+left_f)

        right_f = right_roi_files[group]
        right_rois = jnp.load(path+'/roi_masks/'+right_f)

        area_idx = roi_name_map[area]
        left_rois = jnp.where(left_rois == area_idx)[0]
        right_rois = jnp.where(right_rois == area_idx)[0]
        lh_roi_fmri = jnp.take(left_fmri, left_rois, axis=1)
        rh_roi_fmri = jnp.take(right_fmri, right_rois, axis=1)
        return lh_roi_fmri, rh_roi_fmri
```
