## Supplementary material for "Modelling The Brain’s Response To Natural Scenes In The Bottleneck Space": Source code & results.: downsampler.py

```
import jax.numpy as jnp
import numpy as np

class PCA:
    """
    """
    def __init__(self, dim):
        self.dim = dim

    def fit(self, data):
        data = data.T
        N_dim, N_samples = data.shape
        assert self.dim < N_samples and self.dim < N_dim

        self.μ = jnp.mean(data, axis=1, keepdims=True)
        self.σ = jnp.ones((N_dim, 1))
        data = (data - self.μ) / self.σ

        if N_dim < N_samples:
            C = jnp.einsum("ik,jk->ij", data, data) / (N_samples - 1)
            self.eigenvalues, self.U = jnp.linalg.eigh(C)
            self.eigenvalues = self.eigenvalues[::-1]
            self.U = self.U[:, ::-1]
            self.λ = jnp.sqrt(self.eigenvalues)
            self.U = self.U[:, : self.dim]
        else:
            D = (jnp.einsum("ki,kj->ij", data, data)/ N_dim)
            self.eigenvalues, V = jnp.linalg.eigh(D)
            self.eigenvalues = self.eigenvalues[::-1]
            V = V[:, ::-1]
            self.eigenvalues = self.eigenvalues[: self.dim] * (N_dim / (N_samples - 1))
            self.λ = jnp.sqrt(self.eigenvalues)
            S_inv = (1 / jnp.sqrt(self.eigenvalues * (N_samples - 1)))[jnp.newaxis, :]
            VS_inv = V[:, : self.dim] * S_inv
            self.U = jnp.einsum("ij,jk->ik",data, VS_inv)

        return self

    def transform(self, X):
        X = jnp.asarray(X).T
        return jnp.einsum("ji,jk->ik", self.U, (X - self.μ) / self.σ).T

    def inverse_transform(self, X):
        X = jnp.asarray(X).T
        return (jnp.einsum("ij,jk->ik", self.U, X) * self.σ + self.μ).T
    
    def sample(self, n=1):
        return jnp.array(np.random.normal(size=(self.dim, n)) * np.array(self.λ)[:, np.newaxis]).T
```
