## Supplementary material for "Modelling The Brain’s Response To Natural Scenes In The Bottleneck Space": Source code & results.: evaluation.py

```
import os
import jax.numpy as jnp
import matplotlib.pyplot as plt

def mean_roi_correlation(path,lh_correlation,rh_correlation):
    """
    """
    roi_mapping_files = ['mapping_prf-visualrois.npy', 
                         'mapping_floc-bodies.npy',
                         'mapping_floc-faces.npy', 
                         'mapping_floc-places.npy',
                         'mapping_floc-words.npy', 
                         'mapping_streams.npy']
    
    roi_name_maps = []
    for r in roi_mapping_files:
        roi_name_maps.append(jnp.load(path+'/roi_masks/'+r,allow_pickle=True).item())

    left_roi_files = ['lh.prf-visualrois_challenge_space.npy',
                    'lh.floc-bodies_challenge_space.npy', 
                    'lh.floc-faces_challenge_space.npy',
                    'lh.floc-places_challenge_space.npy', 
                    'lh.floc-words_challenge_space.npy',
                    'lh.streams_challenge_space.npy']

    right_roi_files = ['rh.prf-visualrois_challenge_space.npy',
                    'rh.floc-bodies_challenge_space.npy', 
                    'rh.floc-faces_challenge_space.npy',
                    'rh.floc-places_challenge_space.npy', 
                    'rh.floc-words_challenge_space.npy',
                    'rh.streams_challenge_space.npy']
        
    lh_challenge_rois = [jnp.load(path+'/roi_masks/'+file) for file in left_roi_files]
    rh_challenge_rois = [jnp.load(path+'/roi_masks/'+file) for file in right_roi_files]

    roi_names = []
    lh_roi_correlation = []
    rh_roi_correlation = []
    for r1 in range(len(lh_challenge_rois)):
        for r2 in roi_name_maps[r1].items():
            if r2[0] != 0: 
                roi_names.append(r2[1])
                lh_roi_idx = jnp.where(lh_challenge_rois[r1] == r2[0])[0]
                rh_roi_idx = jnp.where(rh_challenge_rois[r1] == r2[0])[0]
                lh_roi_correlation.append(lh_correlation[lh_roi_idx])
                rh_roi_correlation.append(rh_correlation[rh_roi_idx])

    roi_names.append('All vertices')
    lh_roi_correlation.append(lh_correlation)
    rh_roi_correlation.append(rh_correlation)

    left_roi_correlation = [lh_roi_correlation[r].mean() for r in range(len(lh_roi_correlation))]
    right_roi_correlation = [rh_roi_correlation[r].mean() for r in range(len(rh_roi_correlation))]
    return left_roi_correlation, right_roi_correlation, roi_names
    


def plot_results(left_roi_correlation, right_roi_correlation, roi_names):
    """
    """
    plt.figure(figsize=(18,6))
    x = jnp.arange(len(roi_names))
    width = 0.30
    plt.bar(x - width/2, left_roi_correlation, width, label='Left Hemisphere')
    plt.bar(x + width/2, right_roi_correlation, width,label='Right Hemishpere')
    plt.xlim(left=min(x)-.5, right=max(x)+.5)
    plt.ylim(bottom=0, top=1)
    plt.xlabel('ROIs')
    plt.xticks(ticks=x, labels=roi_names, rotation=60)
    plt.ylabel('Median Pearson\'s $r$')
    plt.legend(frameon=True, loc=1)


def zipdir(path, ziph):
    """
    """
    for root, dirs, files in os.walk(path):
        for file in files:
            ziph.write(os.path.join(root, file), 
                       os.path.relpath(os.path.join(root, file), 
                                       os.path.join(path, '..')))
```
