## Supplementary material for "Modelling The Brain’s Response To Natural Scenes In The Bottleneck Space": Source code & results.: metrics.py

```
import jax 
import jax.numpy as jnp

@jax.jit
def batch_pearsonr(x, y):
    x = jnp.asarray(x).T
    y = jnp.asarray(y).T
    x = x - jnp.expand_dims(x.mean(axis=1), axis=-1)
    y = y - jnp.expand_dims(y.mean(axis=1), axis=-1)
    numerator = jnp.sum(x * y, axis=-1)
    sum_of_squares_x = jnp.einsum('ij,ij -> i', x, x)
    sum_of_squares_y = jnp.einsum('ij,ij -> i', y, y)
    denominator = jnp.sqrt(sum_of_squares_x * sum_of_squares_y)
    return numerator / denominator
```
