## Supplementary material for "Modelling The Brain’s Response To Natural Scenes In The Bottleneck Space": Source code & results.: model.py

```
import jax
import optax
import haiku as hk
import jax.numpy as jnp
import flax.linen as nn

# Model

class Model(nn.Module):
  image_dim: int    
  hidden_expansion: int               
  left_dim: int
  right_dim: int
  dropout: int
  alpha: float
  beta: float

  def setup(self):
     wide_dim = self.image_dim * self.hidden_expansion
     self.widening = nn.Dense(wide_dim)
     self.residual = LinearResidualBlock(wide_dim)
     self.leak_projection = nn.Dense(wide_dim)
     self.left_dense = nn.Dense(self.left_dim)
     self.right_dense = nn.Dense(self.right_dim)

  def __call__(self, x, structured_noise, leak=None):
    x = self.widening(x)
    x = self.residual(x)
    x = Dropout(x, self.dropout)

    structured_noise = self.leak_projection(structured_noise)
    
    if leak is None:
      x = x - ((self.beta + self.alpha) * structured_noise)
    else:
      leak = self.leak_projection(leak)
      x = x + (self.alpha * leak) + (self.beta * structured_noise)

    x /= (1 + self.alpha + self.beta)
    left = self.left_dense(x)
    right = self.right_dense(x)
    return left, right


class LinearResidualBlock(nn.Module): 
  dims: int
  @nn.compact
  def __call__(self, x):
    y = nn.Dense(self.dims)(x)
    y = nn.PReLU()(y)               
    return x + y

@jax.jit
def Dropout(x, p):
   rng = next(hk.PRNGSequence(jax.random.PRNGKey(42)))
   return hk.dropout(rng, p, x)

@jax.jit
def SSE(x, y):
   return jnp.sum((x - y) ** 2, axis=-1)
```
