## Supplementary material for "Modelling The Brain’s Response To Natural Scenes In The Bottleneck Space": Source code & results.: trainer.py

```
import os
import optax
import pickle
import jax.numpy as jnp
from  src.metrics import batch_pearsonr

def train(train_forward, eval_forward, params, optimizer, data, epochs, ckpt_dir, prefix):
  opt_state = optimizer.init(params)
  min_val_corr = float('-inf')
  final_left_corr, final_left_corr = None, None

  for epoch in range(1, epochs+1):
    losses = []
    
    for batch_X, left_Y, right_Y in data.train:
       left_Y = data.left_fmri_downsampler.transform(left_Y)
       right_Y = data.right_fmri_downsampler.transform(right_Y)
       sampled_left = data.left_fmri_downsampler.sample(batch_X.shape[0])
       sampled_right = data.right_fmri_downsampler.sample(batch_X.shape[0])
       structured_noise = jnp.concatenate((sampled_left, sampled_right), axis=-1)
       loss, grads = train_forward(params, batch_X, structured_noise, left_Y, right_Y)
       updates, opt_state = optimizer.update(grads, opt_state, params=params)
       params =  optax.apply_updates(params=params, updates=updates)
       losses.append(loss)
       
    train_loss = jnp.array(losses).mean()
    print("Epoch: {}".format(epoch))
    print("Train loss: {}".format(train_loss))
    left_corr, right_corr, val_corr = evaluate(eval_forward, params, data)

    if val_corr > min_val_corr:
      print('Saving Parameters...')
      pickle.dump(params, open(os.path.join(ckpt_dir, prefix), "wb"))
      min_val_corr = val_corr.copy()
      final_left_corr = left_corr.copy()
      final_right_corr = right_corr.copy()

    print("\n")
    
  return params, final_left_corr, final_right_corr


def evaluate(eval_forward, params, data):
  X, true_left, true_right = data.val
  downsampled_left = data.left_fmri_downsampler.transform(true_left)
  downsampled_right = data.right_fmri_downsampler.transform(true_right)

  sampled_left = data.left_fmri_downsampler.sample(X.shape[0])
  sampled_right = data.right_fmri_downsampler.sample(X.shape[0])
  structured_noise = jnp.concatenate((sampled_left, sampled_right), axis=-1)

  (low_dim_left_corr, 
    low_dim_right_corr, 
    left_preds, 
    right_preds) = eval_forward(params, 
                                X, 
                                structured_noise,
                                downsampled_left, 
                                downsampled_right)
  

  left_preds = data.left_fmri_downsampler.inverse_transform(left_preds)
  right_preds = data.right_fmri_downsampler.inverse_transform(right_preds)
  left_corr = batch_pearsonr(true_left, left_preds)
  right_corr = batch_pearsonr(true_right, right_preds)

  reconstructed_left = data.left_fmri_downsampler.inverse_transform(downsampled_left)
  reconstructed_right = data.right_fmri_downsampler.inverse_transform(downsampled_right)
  left_reconstruction_corr = batch_pearsonr(true_left, reconstructed_left).mean()
  right_reconstruction_corr = batch_pearsonr(true_right, reconstructed_right).mean()
  val_corr = jnp.concatenate((left_corr, right_corr)).mean()

  print("Low Dim Left Mean Correlation: {}".format(low_dim_left_corr.mean()))
  print("Low Dim Right Mean Correlation: {}".format(low_dim_right_corr.mean()))
  print("Left Reconstruction Mean Correlation: {}".format(left_reconstruction_corr))
  print("Right Reconstruction Mean Correlation: {}".format(right_reconstruction_corr))
  print("Final Left Mean Correlation: {}".format(left_corr.mean()))
  print("Final Right Mean Correlation: {}".format(right_corr.mean()))
  print('Total Correlation:', val_corr)
  return left_corr, right_corr, val_corr
```
