## Supplementary figures and images for "Modelling The Brain’s Response To Natural Scenes In The Bottleneck Space"

### bottleneck.png

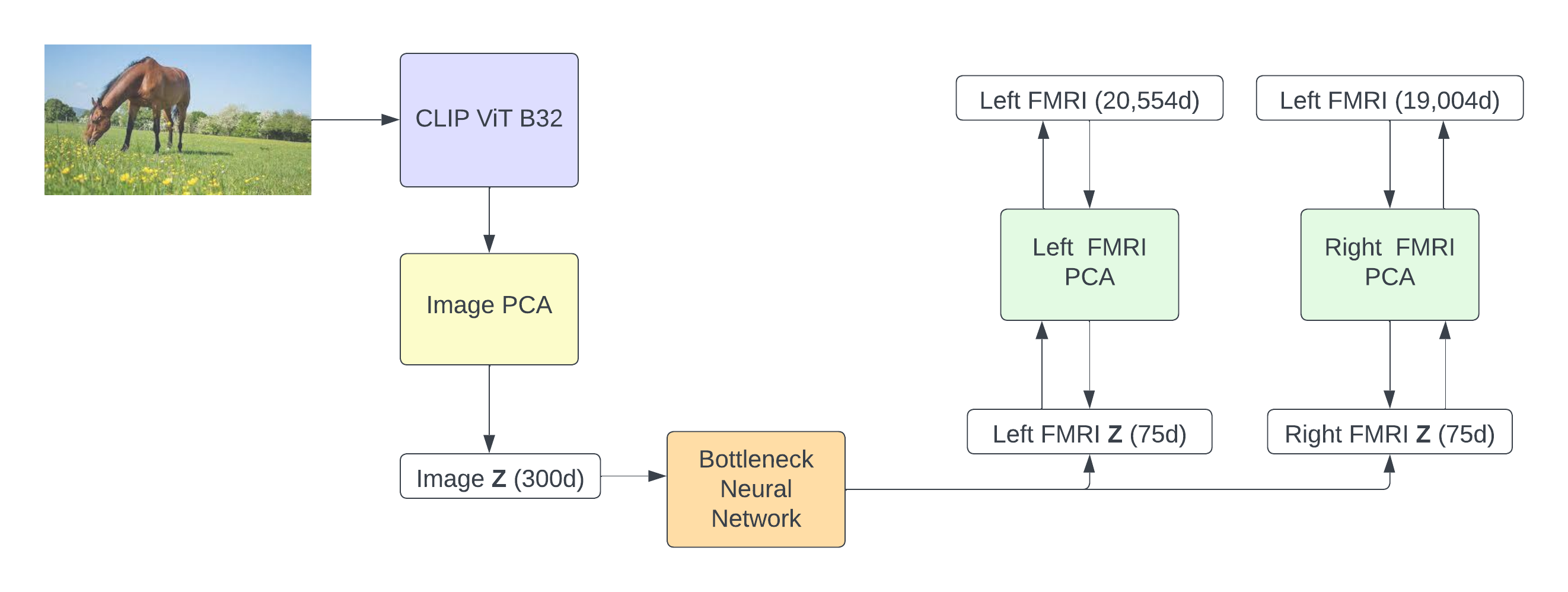

### equations.png

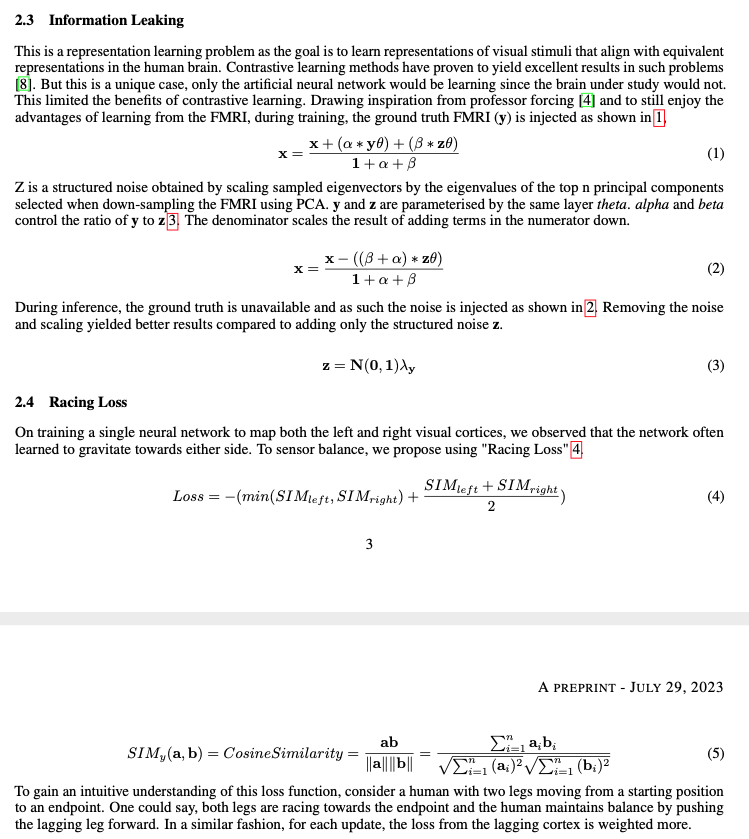

### network.png

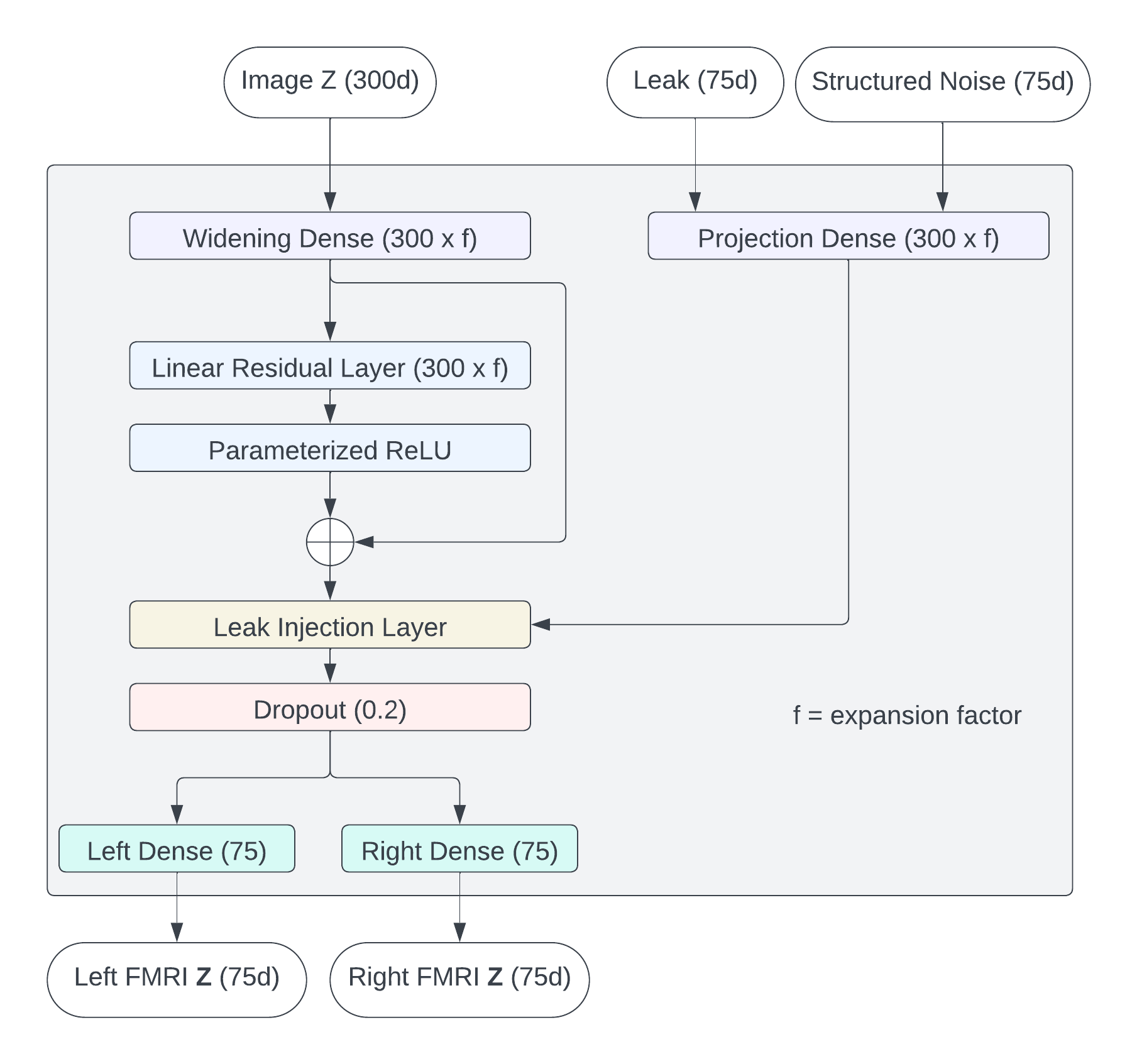

### scores.png

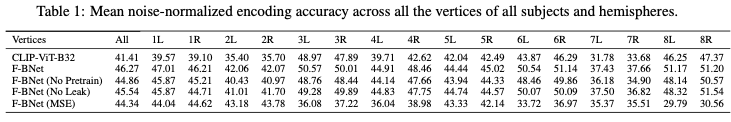
